# scCompare: a web app for single-cell RNA sequencing dataset comparisons across multiple auto-immune diseases

**DOI:** 10.1101/2025.02.04.636484

**Authors:** Shweta Yadav, Pranav Sahasrabudhe, Mulini Pingili, Dan Chang, Jing Wang

**Author notes:** **Corresponding author:** Jing Wang, PhD, AbbVie Inc, 200 Sidney St, Cambridge, MA 02139, USA. **Author Contributions** Conceptualization: JW; Interface and database development: SY, PS, MP; Data curation: SY; Project Administration: JW, DC; Wrote the draft: JW, DC, SY. All authors critically reviewed the manuscript, provided feedback, and approved the manuscript. **Disclosures** JW, DC, and PS are employees of AbbVie. SY and MP are contractors for AbbVie. The design, study conduct, and financial support for this research were provided by AbbVie. AbbVie participated in the interpretation of data, review, and approval of the publication.

## Abstract

**Motivation:** Single-cell RNA sequencing (scRNA-seq) datasets have been widely used to identify the cell types and marker genes with pivotal roles in driving pathogenesis and progression of auto-immune diseases. Comparative analysis of different cell types or diseases across multiple scRNA-seq datasets can reveal the homogeneous and heterogeneous pathogenesis, and a comprehensive web-based comparison tool that could streamline this process is not yet available.

**Results:** We introduce scCompare, a web-based platform for scRNA-seq comparisons in autoimmune diseases. scCompare includes 2,125 differential gene lists from 100 scRNA-seq datasets in 22 auto-immune diseases and a query system supporting 170 standardized keywords from four attributes (disease, cell type, tissue, and treatment). scCompare also provides three modules enabling several comparative analysis and visualization options. geneQuery supports comparisons of queried genes across differential gene lists identified from multiple scRNA-seq datasets. DEGEnricher performs cell-type-specific enrichment analysis across studies based on a user-input gene list. DEGCompare allows interactive comparisons of multiple differential gene lists of many studies and performs pathway enrichment analyses. Using two case studies as examples, we demonstrated that scCompare represents a unique platform for biologists to identify, compare and validate the pathogenesis at the single-cell levels among auto-immune diseases.

**Availability and implementation:** scCompare is freely available at https://sccompare.shinyapps.io/main/. The source code is available at https://github.com/abbviegrc/scCompare.

## Introduction

Autoimmune diseases occur when the immune system produces antibodies to attack the body’s own cells(Smith and Germolec, 1999). Over 80 types of autoimmune diseases have been identified, which affect nearly 4% of the world’s population with a steady rise over the last decades(Bragazzi, et al., 2017). Although many drugs have been developed to treat different autoimmune diseases (e.g. non-steroidal anti-inflammatory drugs, glucocorticoids, and biologic drugs)(Li, et al., 2017), there is still a significant unmet need for new targeted therapies for inadequate-response patients(Reves, et al., 2021).

Single-cell RNA sequencing (scRNA-seq) technology has become the state-of-the-art approach for understanding the pathogenesis of autoimmune diseases by unravelling the heterogeneity and complexity of RNA transcripts within individual cells(Yang, et al., 2023), which can provide important implications for novel therapies (Alivernini, et al., 2020). Several platforms have been developed to host, share, and explore the scRNA-seq datasets (e.g. Broad Institute single cell portal (https://singlecell.broadinstitute.org/single_cell), single-cell expression atlas(Moreno, et al., 2021), and Bioturing (https://bioturing.com/)). After selecting one scRNA-seq dataset, these platforms can visualize the expression of queried genes across cell clusters, identify the differentially expressed genes (DEGs) among cell clusters or between conditions within one cell cluster, and perform the more comprehensive analyses (e.g. trajectory analysis). With the exponential growth of scRNA-seq datasets, many studies have performed integration of multiple scRNA-seq datasets to identify shared or heterogenous pathogenesis across diseases (Korsunsky, et al., 2022). However, there is no web-based platform that can easily perform this type of analysis. Although the Broad Institute single cell portal can query multiple genes across multiple single-cell studies, it cannot filter the studies by disease, tissue or other keywords, nor provide the significance of genes across studies. scIBD (Nie, et al., 2023) is a data portal for CD and UC that integrated 1.13 million cells from 12 public datasets. However, all results must be regenerated when including more datasets. Although DISCO (Li, et al., 2022) and Bioturing allow customized integration by selecting samples or datasets, it is difficult to build a cell atlas and manually perform all down-stream comparisons due to the complexity and massive time consumption of data integration.

To streamline the interactive analysis of scRNA-seq comparisons, we developed scCompare, a web-based comparison tool for auto-immune diseases. scCompare has five key features: (i) 2,125 differentially expressed genes (DEGs) from 100 scRNA-seq datasets in 22 auto-immune diseases. (ii) a query system for scRNA-seq datasets and differential gene lists supporting 170 keywords from four categories. (iii) geneQuery: dataset comparisons across DEGs based on the queried genes. (v) DEGEnricher: identification of the enriched DEGs based on a user-input gene list. (v) DEGCompare: interactive comparisons of DEGs generated from multiple scRNA-seq datasets and identification of enriched pathways based on the comparison result. We also provided two case studies to demonstrate the utility of this unique tool.

## Materials and Methods

### scRNA-seq collection and metadata curation

22 auto-immune diseases were searched in Bioturing (https://bioturing.com/) to retrieve 100 scRNA-seq datasets (Figure 1 and supplementary Table S1). For each dataset, DEGs between cells from one cell type with at least 100 cells and all other cells was identified based on venice method. All genes were included (no threshold filter) except those expressed in less than 10% of cells in two comparison groups. If the dataset included the control/healthy samples, the DEGs between cells from disease samples and control samples were also identified for each cell type.

**Figure 1.**
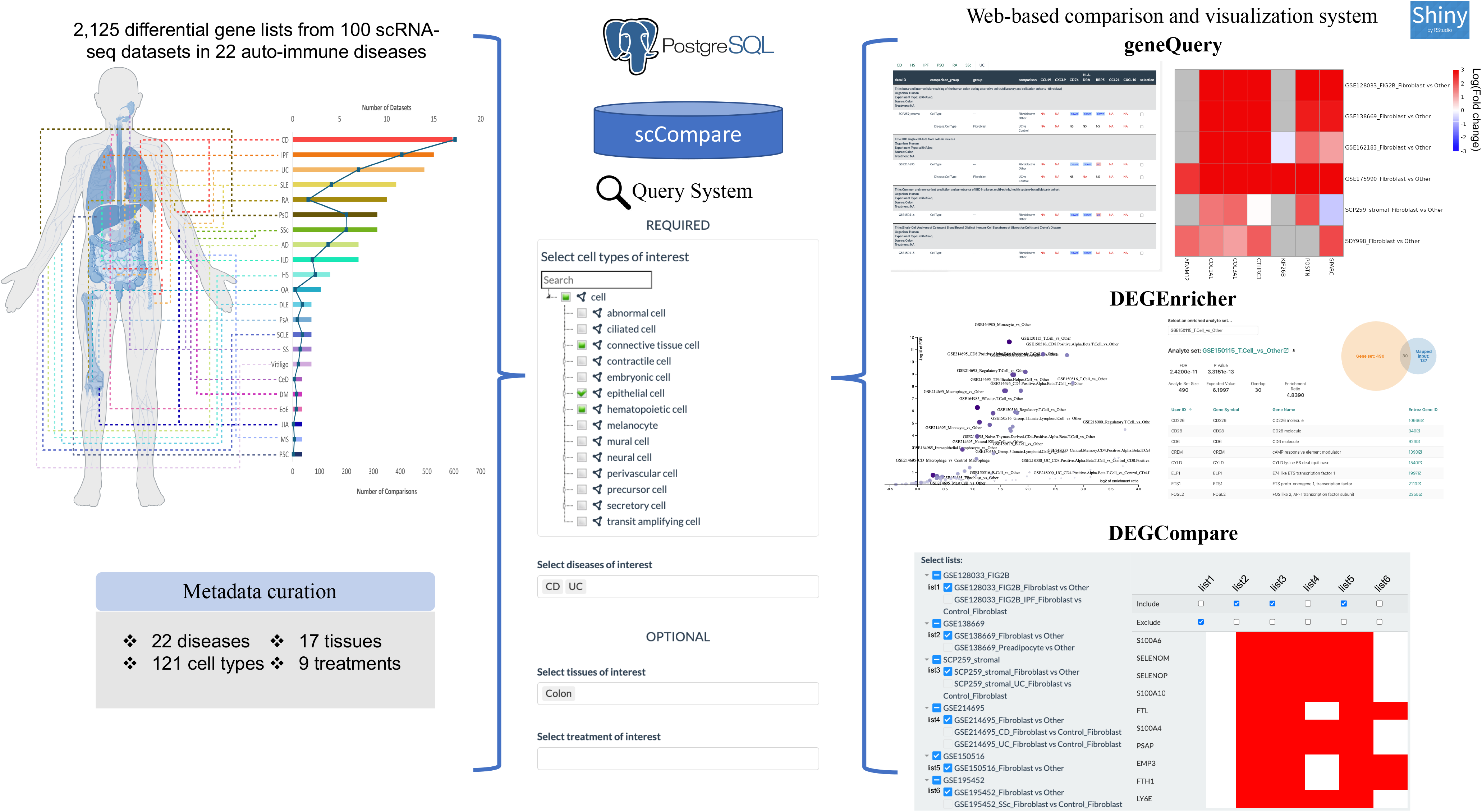
scCompare for identifying, comparing, and validating the pathogenesis among auto-immune diseases. Left panel: 2,125 DEGs from 100 scRNA-seq datasets in 22 auto-immune diseases. Bar plot and line plot represented the number of datasets and comparisons for each disease, respectively. The dot line represented the sources of the scRNA-seq datasets from each disease. Middle panel: scCompare query system based on 170 keywords from four attributes. Right panel: three comparison modules.

For the longitudinal treatment datasets, the DEGs between cells from different timepoints were identified for each cell type. In total, scCompare included 2,125 differential comparisons from 100 scRNA-seq studies (Figure 1 and supplementary Table S2). The metadata were manually curated based on the 170 keywords we defined from four attributes: 23 diseases/control, 121 cell types, 17 tissues, and 9 treatments (supplementary Table S3). The cell types were organized based on cell ontology (Diehl, et al., 2016), while diseases/control, tissues, and treatments were mapped to MedDRA LLT (Medical Dictionary for Regulatory Activities Lowest Level Term)(Brown, et al., 1999) and NCIT (National Cancer Institute Thesaurus) (Fragoso, et al., 2004). All comparison results and metadata were stored in a PostgreSQL database.

## Data analysis modules

scCompare allows accurate data and DEGs selection based on the user query from four attributes (e.g. fibroblast cells vs others from CD and UC datasets). After selecting the DEGs, three modules are available for users to compare the results (Figure 1).

### geneQuery

geneQuery module enables easy comparison of significance of queried genes (less than 10) in multiple DEGs. After selecting multiple DEGs, geneQuery can also plot the heatmap based on log(Fold change) or signed −log(p-value).

## DEGEnricher

For a long list of queried genes (greater than 10), scCompare provides DEGEnricher module to perform enrichment analysis of the gene lists against the selected DEGs by WebGestaltR package(Liao, et al., 2019). Multiple interactive plots provided by WebGestaltR are imbedded in DEGEnricher for visualization of enrichment results (e.g. bar plots and volcano plots for result summary).

## EDEGCompare

DEGCompare module provides an interactive table to compare multiple DEGs. The DEGs can be from the same datasets (e.g. marker genes from different cells) or different datasets (e.g. marker genes of one cell type from different diseases). Checking “Include” or “Exclude” boxes under each DEG, the filtered genes will be shown in a table, which is a more powerful comparison visualization tool than Venn diagram for multiple gene lists (Figure 1). DEGCompare can also perform Reactome pathway enrichment analysis for the filtered genes by WebGestaltR package(Liao, et al., 2019), or plot Venn diagram if the number of selected DEGs were less or equal than four.

## Case studies

### Evaluating the shared fibroblast states across seven auto-immune diseases

Korsunsky et al. (Korsunsky, et al., 2022) identified SPARC^+^COL3A1^+^ and CXCL10^+^CCL19^+^ fibroblasts shared by multiple auto-immune diseases, which can be estimated by multiple scRNA-seq datasets and expanded to more diseases in the geneQuery module. SPARC^+^COL3A1^+^ fibroblast includes eight representative marker genes while CXCL10^+^CCL19^+^ fibroblast includes seven representative markers (supplementary Table S4). Based on the following keywords: cell type keyword (fibroblast) and disease keywords (CD (crohn’s disease), HS (hidradenitis suppurativa), IPF (idiopathic pulmonary fibrosis), PsO (psoriasis), RA (rheumatoid arthritis), SSc (systemic scleroderma), and UC (ulcerative colitis)), seven DEGs between fibroblast cells and other cells from seven scRNA-seq datasets were selected to query the SPARC^+^COL3A1^+^ and CXCL10^+^CCL19^+^ marker genes. As shown in supplementary Figure S1A, SPARC^+^COL3A1^+^ marker genes had higher expression in fibroblasts than other cells in most of seven diseases. However, most of CXCL10^+^CCL19^+^ marker genes were not included in seven DEGs due to no expression in at least 90% of fibroblast cells (supplementary Figure S1B and supplementary Figure S2), which was confirmed in more scRNA-seq datasets from seven diseases (supplementary Figure S3). CD74 and HLA-DRA had significantly lower expression in fibroblasts from seven diseases (supplementary Figure S1B), which was consistent with UMAP plots of seven selected scRNA-seq datasets from Bioturing (supplementary Figure S2). Thus, only SPARC^+^COL3A1^+^ fibroblast may represent the shared fibroblast across seven diseases. Together, this demonstrates scCompare’s capacity for identifying the similar or heterogeneous cell types across diseases.

### Integrating GWAS genes of inflammatory bowel disease with scRNA-seq datasets to identify the pathogenic cell types

Integrating genome-wide association studies (GWAS) with scRNA-seq dataset can infer the cell types by which genetic variants influence diseases (Jagadeesh, et al., 2022), and this analysis can be implemented by the DEGEnricher module. For inflammatory bowel disease (IBD), 152 GWAS genes with L2G score>0.5 were downloaded from (Barrio-Hernandez, et al., 2023). Based on the following keywords: cell type keywords (“fibroblast”, “epithelial cell”, “leukocyte”), disease keywords (“CD”, “UC”), and tissue keyword (“Colon”), 226 DEGs from 12 scRNA-seq datasets were selected and significant gene lists were identified under FDR<0.05 and log(Fold change)>0.25. As shown in supplementary Table S4, 73 DEG lists were enriched with IBD GWAS genes under FDR<0.05. Seven out of top 10 enriched DEG lists were related to T cells, which was consistent with previous IBD cis-eQTL results (Hu, et al., 2021). Overall, this exemplifies the utility of scCompare in identifying GWAS-related cell types in scRNA-seq datasets.

## Discussions

Because of a lack of web-based tools to facilitate multiple scRNA-seq comparisons, we present scCompare, a manually curated scRNA-seq comparison tool with comprehensive query system and interactive data comparison and analysis functions. scCompare has the following unique features compared to the existing web-based scRNA-seq tools. First, scCompare includes 2,125 differential comparisons from 100 scRNA-seq datasets in 22 auto-immune diseases and a query system supporting 170 keywords from four attributes. Second, geneQuery supports comparisons of queried genes (less than 10 genes) across multiple DEGs. Third, DEGEnricher identifies the enriched DEGs based on a user-input gene list (over 10 genes). Fourth, DEGCompare module allows interactive comparisons of DEGs and perform pathway enrichment analyses based on the comparison results. Because of the time and memory consumption, scCompare does not provide visualization functions (e.g. UMAP) for multiple scRNA-seq datasets. scCompare is a powerful tool to enable users to evaluate genes of interest across diseases, cell types and studies to identify relevant signals, and users can then leverage Broad Institute single cell portal or Bioturing to take a deep dive into individual scRNA-seq datasets. In the future, we will continue to incorporate new datasets as well as novel functions to enhance scCompare.

## Supporting information

Tutorial

## Reference

Alivernini, S., et al. Distinct synovial tissue macrophage subsets regulate inflammation and remission in rheumatoid arthritis. Nat Med 2020;26(8):1295–1306.

Barrio-Hernandez, I., et al. Network expansion of genetic associations defines a pleiotropy map of human cell biology. Nat Genet 2023;55(3):389–398.

Bragazzi, N.L., et al. Public health awareness of autoimmune diseases after the death of a celebrity. Clin Rheumatol 2017;36(8):1911–1917.

Brown, E.G., Wood, L. and Wood, S. The medical dictionary for regulatory activities (MedDRA). Drug Saf 1999;20(2):109–117.

Diehl, A.D., et al. The Cell Ontology 2016: enhanced content, modularization, and ontology interoperability. J Biomed Semantics 2016;7(1):44.

Fragoso, G., et al. Overview and utilization of the NCI thesaurus. Comp Funct Genomics 2004;5(8):648–654.

Hu, S., et al. Inflammation status modulates the effect of host genetic variation on intestinal gene expression in inflammatory bowel disease. Nat Commun 2021;12(1):1122.

Jagadeesh, K.A., et al. Identifying disease-critical cell types and cellular processes by integrating single-cell RNA-sequencing and human genetics. Nat Genet 2022;54(10):1479–1492.

Korsunsky, I., et al. Cross-tissue, single-cell stromal atlas identifies shared pathological fibroblast phenotypes in four chronic inflammatory diseases. Med 2022;3(7):481–518 e414.

Li, M., et al. DISCO: a database of Deeply Integrated human Single-Cell Omics data. Nucleic Acids Res 2022;50(D1):D596–D602.

Li, P., Zheng, Y. and Chen, X. Drugs for Autoimmune Inflammatory Diseases: From Small Molecule Compounds to Anti-TNF Biologics. Front Pharmacol 2017;8:460.

Liao, Y., et al. WebGestalt 2019: gene set analysis toolkit with revamped UIs and APIs. Nucleic Acids Res 2019;47(W1):W199–W205.

Moreno, P., et al. User-friendly, scalable tools and workflows for single-cell RNA-seq analysis. Nat Methods 2021;18(4):327–328.

Nie, H., et al. Single-cell meta-analysis of inflammatory bowel disease with scIBD. Nat Comput Sci 2023;3(6):522–531.

Reves, J., Ungaro, R.C. and Torres, J. Unmet needs in inflammatory bowel disease. Curr Res Pharmacol Drug Discov 2021;2:100070.

Smith, D.A. and Germolec, D.R. Introduction to immunology and autoimmunity. Environ Health Perspect 1999;107 Suppl 5:661–665.

Yang, X., et al. Research progress on the application of single-cell sequencing in autoimmune diseases. Genes Immun 2023;24(5):220–235.

