## Supplementary Figures for "scCompare: a web app for single-cell RNA sequencing dataset comparisons across multiple auto-immune diseases"

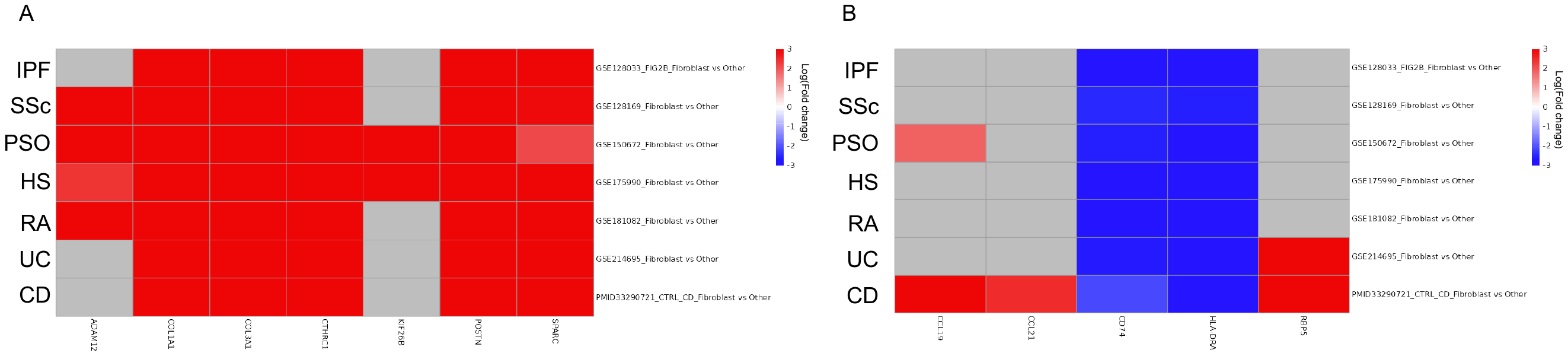

Supplementary Figure S1. Estimation of SPARC^+^COL3A1^+^ and CXCL10^+^CCL19^+^ fibroblasts across seven auto-immune diseases. (A-B) log(Fold change) of seven SPARC^+^COL3A1^+^ marker genes and five CXCL10^+^CCL19^+^ marker genes in seven diseases.

Supplementary Figure S2. Expression of CD74, HLA-DRA, and CXCL10 in seven selected scRNA-seq datasets. CD74 and HLA-DRA had significantly lower expression in fibroblast cells than other cells while CXCL10 did not express in at least 90% of fibroblast cells.

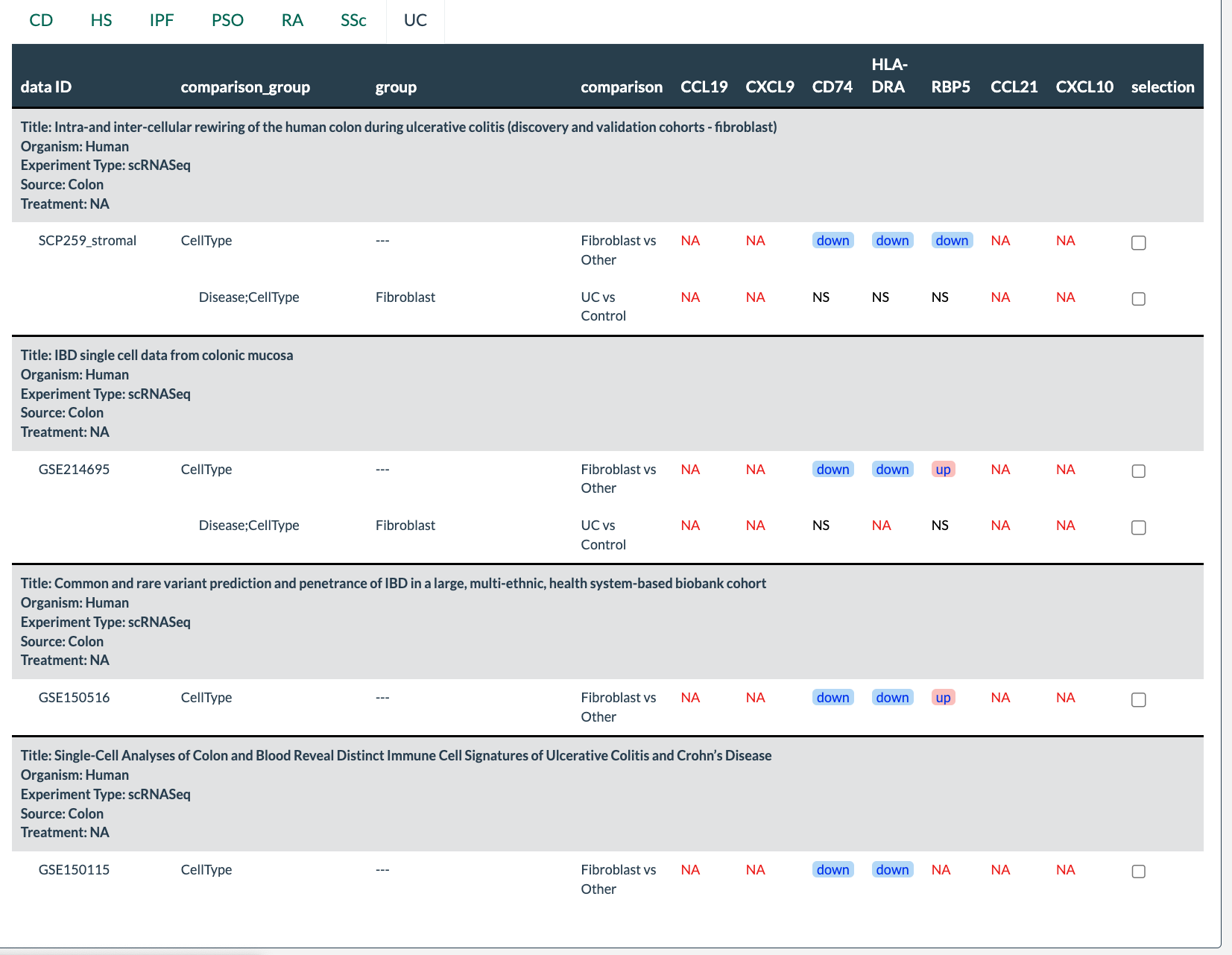

Supplementary Figure S3. Validation of CXCL10^+^CCL19^+^ marker genes in all UC scRNA-seq datasets
