## Supplementary material for "scCompare: a web app for single-cell RNA sequencing dataset comparisons across multiple auto-immune diseases": Tutorial

### User Manual for scCompare

scCompare is a web-based platform for scRNA-seq comparisons in autoimmune diseases.

To begin with the scCompare tool, the user can select the cell type (orange box in Figure 1). Users can either type the cell type of interest in the search box or click on triangle to expand the drop-down list that include 121 cell types. Users are allowed to select multiple cell types. Then, the users are required to select disease of interest (blue box in Figure 1). Users can select one or multiple diseases of interest by typing in the disease name in input box or by clicking on the disease from the drop-down menu. This menu is updated based on the selection made in the cell type section.

After selecting cell type (fibroblast) and disease (AD, CD and PSO), users can select other optional attributes (e.g. tissue or treatment of interest) highlighted in yellow box in Figure 1. Keep in mind that dropdown options change based on the selection made in the previous step.

After confirming selections (green box in Figure 1), related datasets will be displayed in the main console, users can click on checkboxes to select datasets. Users can also click on the dataset_acc (black box in Figure 1) to view the summary of a particular dataset (shown in Figure 2).

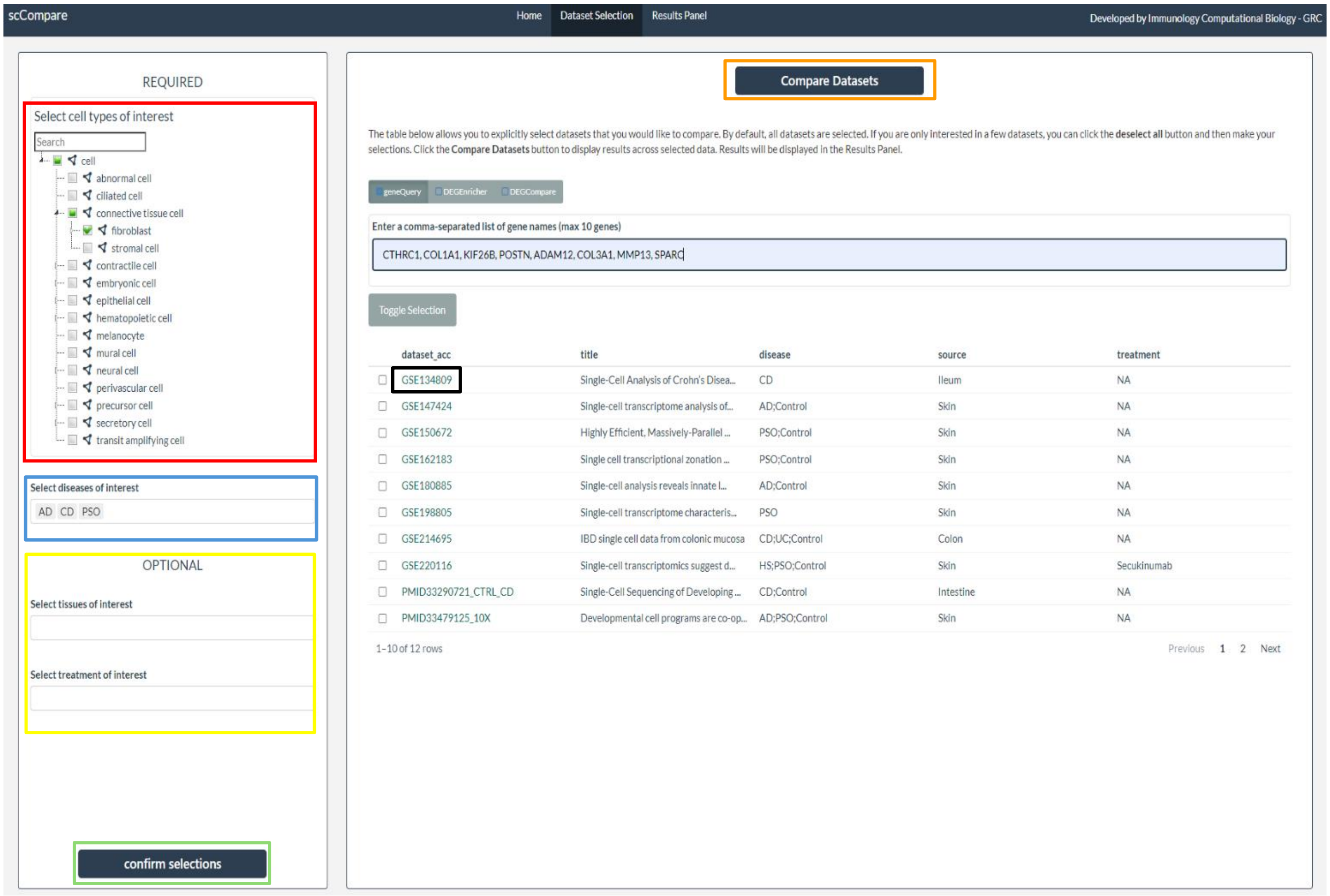


Figure1: Data Selection in scCompare

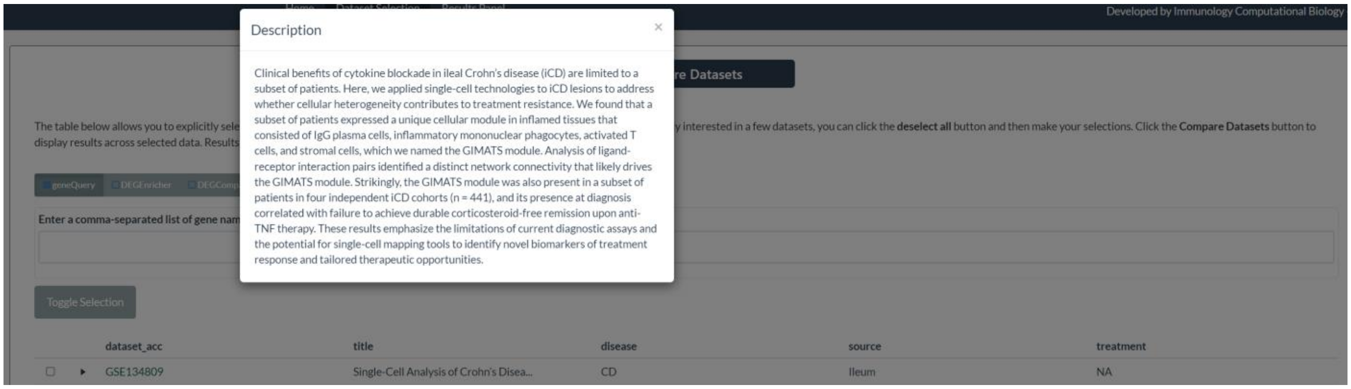


Figure 2: Data Description after clicking on dataset_acc

Now, there are three analysis type options available – **geneQuery, DEGEnricher, DEGCompare** (brown box in Figure 3).

#### 1.1 geneQuery

Users need to enter a list of comma-separated genes, a maximum of 10 genes are allowed.

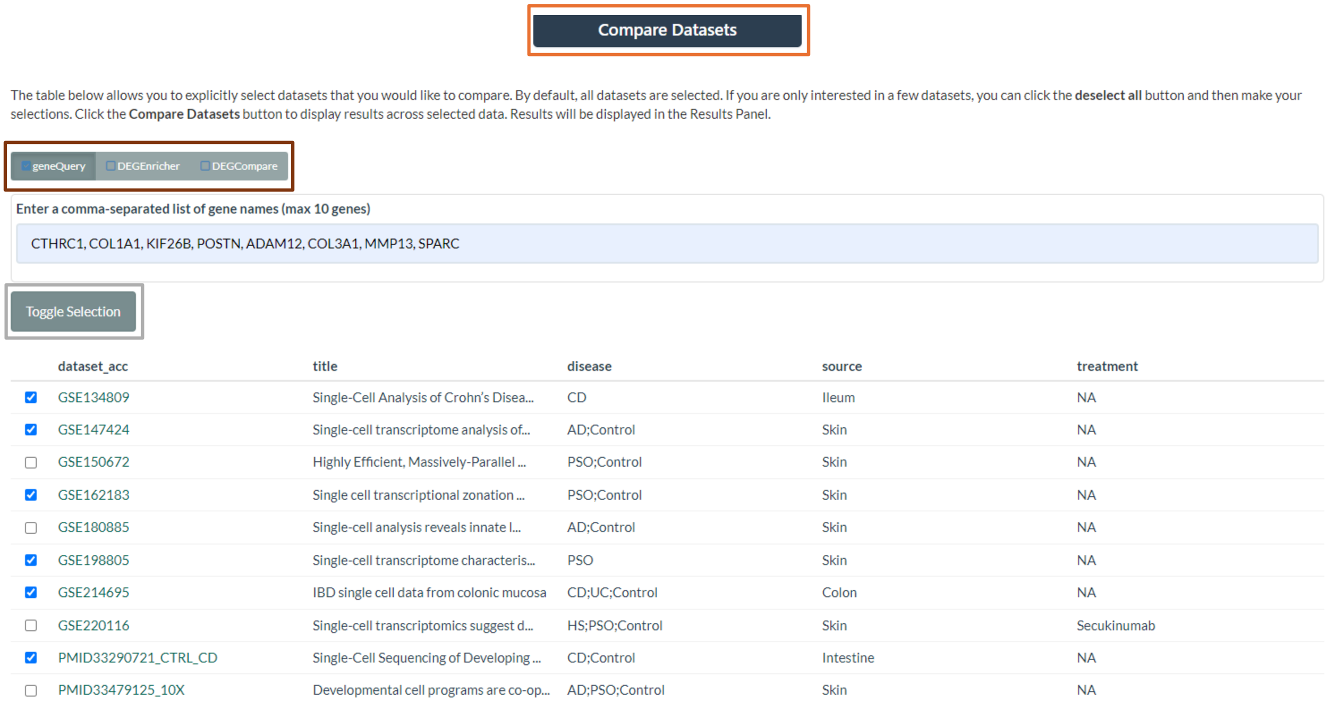


Figure 3: Matching Datasets based on user provided parameters

Next step is to select datasets, users can do so by clicking on the checkboxes or using Toggle Selection button (grey box in Figure 3) to select/deselect all the datasets.

Click on “Compare Datasets” button (orange box in Figure 3).

#### 1.2 DEGEnricher

DEGEnricher performs Enrichment Analysis, users are required to enter their gene list comprising of at least 10 genes (brown box Figure 4). Users can specify Geneset Threshold or Pathway Threshold parameters (blue box in Figure 4) and select the datasets for enrichment analysis. Click on “Compare Datasets” button (orange box in Figure 4) to view results.

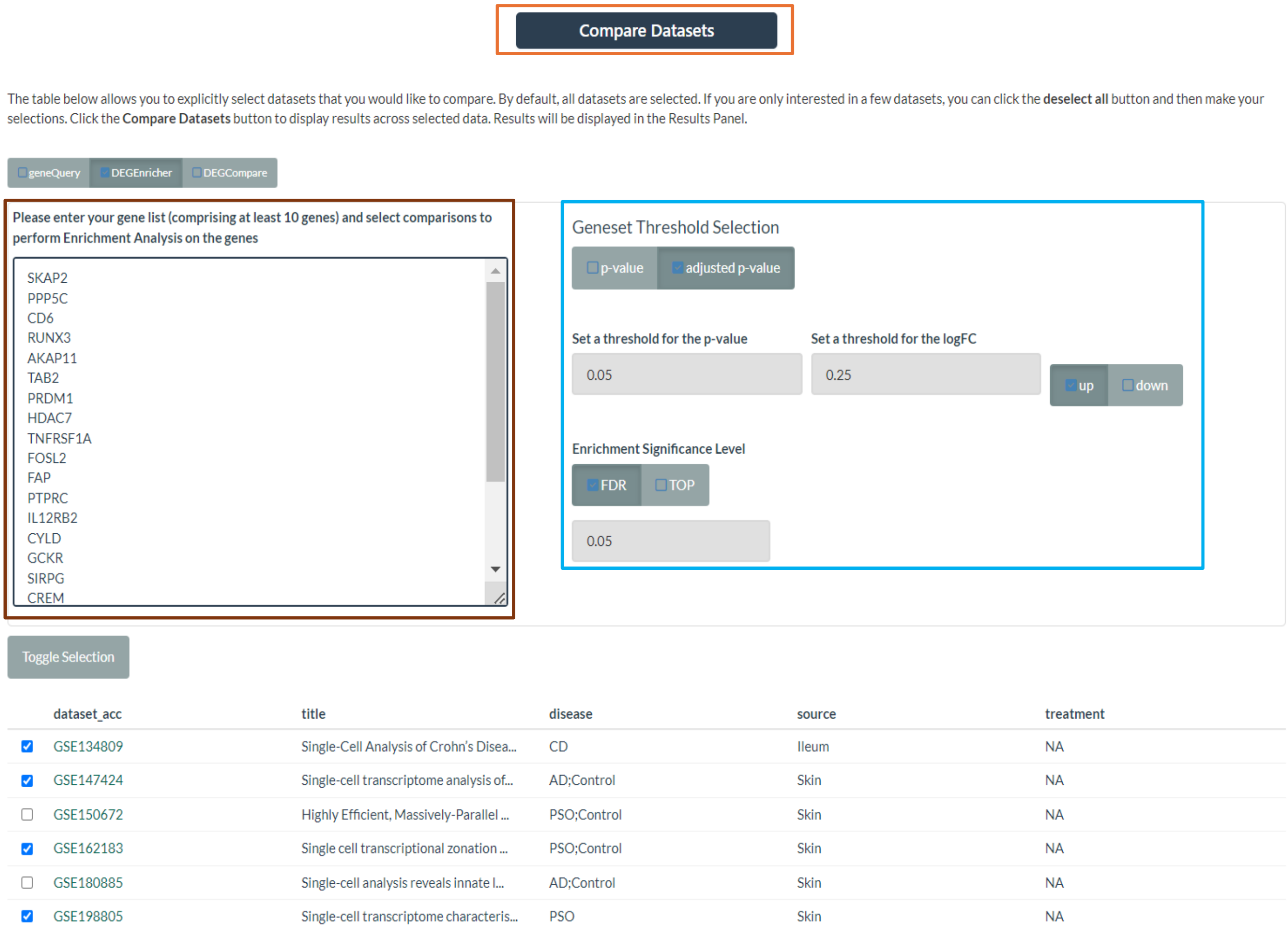

Figure 4: DEGEnricher Tab in scCompare

#### 1.3 DEGCompare

“DEGCompare” allows to compare DEGs and perform pathway enrichment analysis. Users need to specify the threshold for p-value or log fold change (green box in Figure 5) to filter the DEGs and then select datasets of interest. Click on “Compare Datasets” button (orange box in Figure 5) to view results.

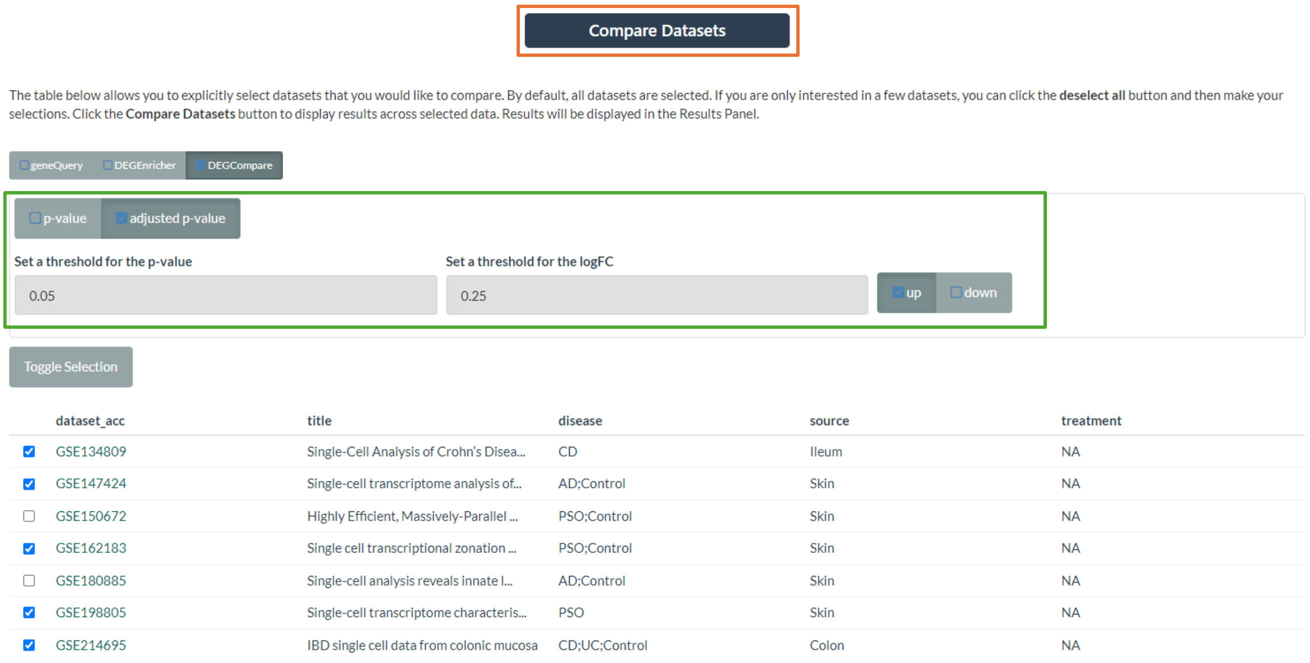


Figure 5: DEGCompare tab in scCompare

#### 2. Results

The comparison results are shown in the "Results Panel" tab.

##### 2.1 geneQuery Result

For geneQuery, there are two sub tabs: Comparisons Table and Expression Visuals. In the "Comparison Tables (Results)" tab, "Summary of Query and Matching Datasets" panel includes query parameters and number of datasets related to each query gene based on each disease (red box in Figure 6). Results panel includes the comparison results with sub-tabs for each disease. The table includes the description of each dataset and comparison and whether the genes of interest are significant in the comparison.

"NS" means gene of interest is not significant in the comparison while red and blue "info" button represent the gene of interest is significantly up- or down-regulated in the comparison. Hovering over the "info" button can get the detailed statistic result. The directionality of the comparison can be found in the "comparison" column. For example, "Connective Tissue vs Others" means up-regulated gene has higher expression in Connective Tissue compared with the other cells.

User can modify the threshold to change the status of each gene in each comparison by clicking the "Table Options" button (orange box in Figure 6).

Users can select one or multiple comparisons for each disease and click on "Visualize Comparison" button to plot the heatmap for each gene in each comparison in the "Expression Visuals" panel.

Please keep in mind that if a dataset has multiple diseases, some comparisons are common across the diseases. For example, in PMID33479125_10X, Connective Tissue Cell vs Other will be displayed under both the tabs- AD and PSO. Users can select this comparison under either one (AD or PSO) or both tabs to generate the heatmap.

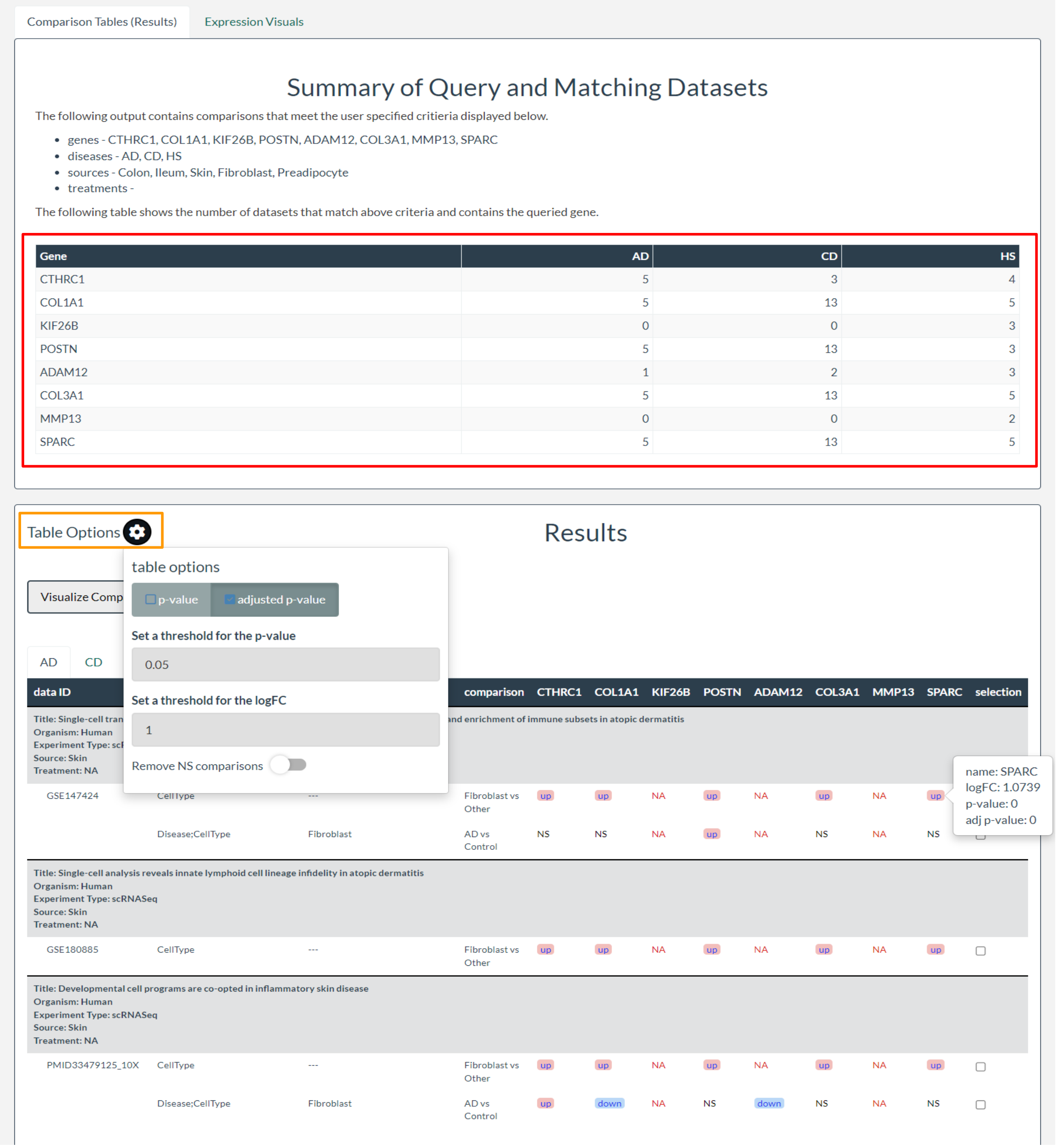

Figure 6: Comparison Tables Results in scCompare

Under Expression Visuals Tab, users can plot the heatmap based on log (Fold change) or signed -log(p-value). Users can also modify the scale for the heatmap plot using the Select ceiling/floor option.

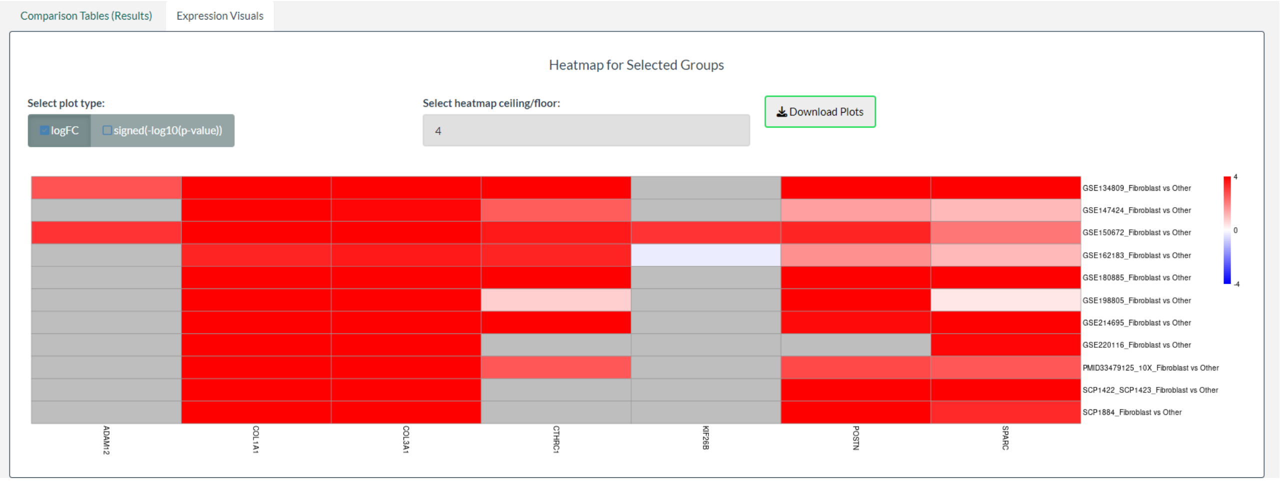


Figure 7: Heatmap under Expression Visuals Tab in scCompare

#### 2.2 DEGEnricher

After entering the gene list and clicking on "Compare Datasets", user is redirected to "Enrichment Analysis" Tab under Results Panel. It displays the enrichment analysis report identifying enriched differential gene lists based on user provided gene list input.

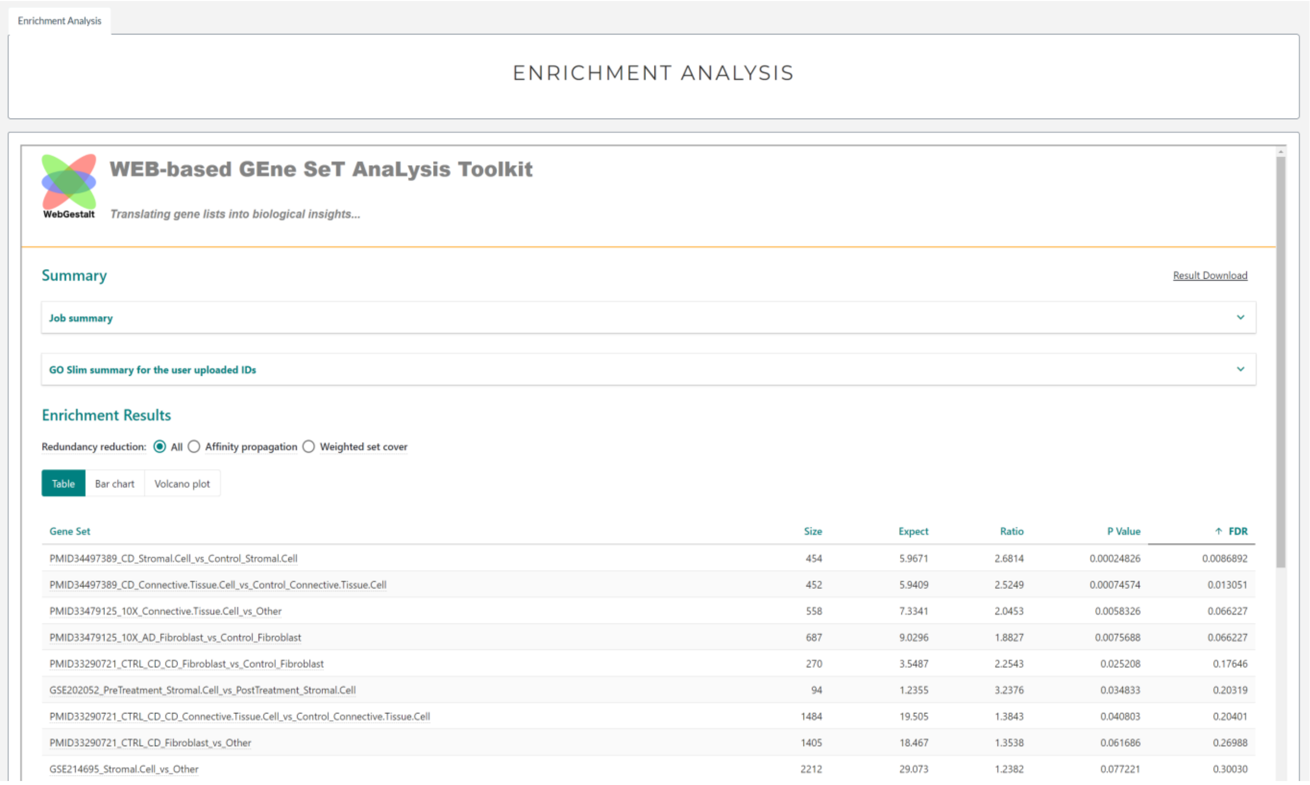


Figure 8: Enrichment results using WebgestaltR

##### 2.3 DEGCompare

DEGCompare allows interactive comparisons of multiple differential gene lists and performs pathway enrichment analyses. Users can select the gene lists from the left-hand side menu (blue box in Figure 9), maximum 7 gene lists can be selected at once.

While comparing the genes among these selected gene lists, users can click on the checkboxes to include or exclude a particular gene list from the visualization (orange box in Figure 9). Users can also click on Enrichment Analysis button (green box in figure 9) to perform pathway enrichment analysis on the selected genes.

"Download Full Results" button (black box in figure 9) allows users to download complete list of genes based on selections made in include/exclude option.

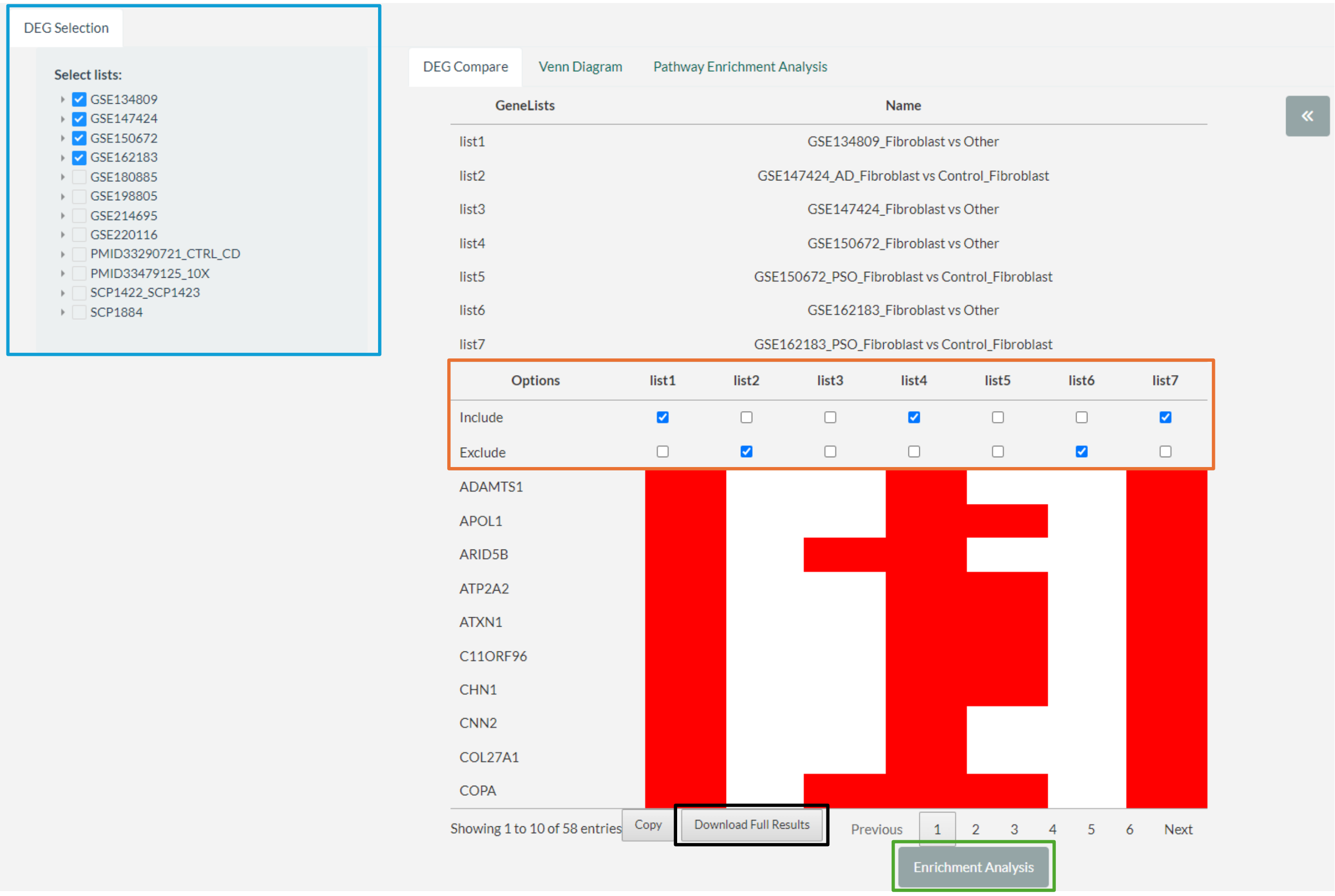


Figure 9: DEGCompare tab under Results Panel

Under "Venn Diagram" tab, by default 4 gene lists are selected. A maximum of 4 gene lists can be selected at once. It provides a Venn plot to identify the number of common genes between the lists. Users can select which four lists they would like to compare from the dropdown menu (black box in Figure 10). Users can also relabel these gene lists in the text box (purple box in Figure 10) below each gene list name.

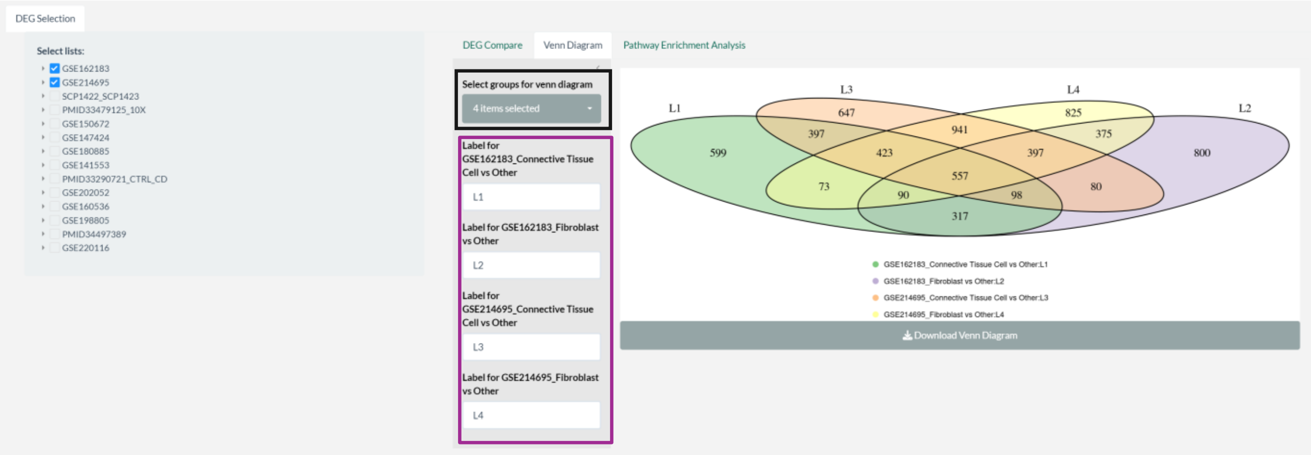

Figure 10: Venn Diagram tab under Results Panel

After clicking on "Enrichment Analysis", User is redirected to Pathway Enrichment Analysis tab, where enrichment report is generated using WebGestaltR. Reactome Pathway enrichment analysis is performed for the filtered genes from DEGCompare.

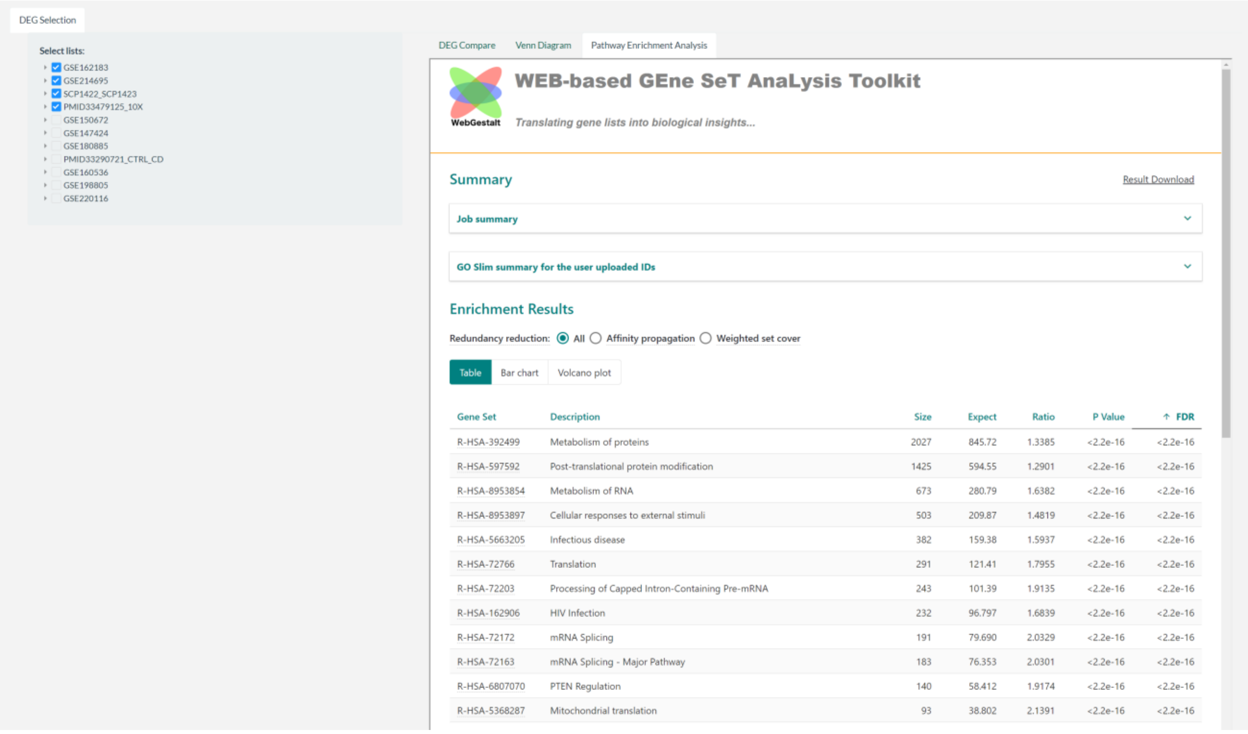


| Gene Set | Description | Size | Expect | Ratio | P Value | FDR |
|---|---|---|---|---|---|---|
| R-HSA-392499 | Metabolism of proteins | 2027 | 845.72 | 1.3385 | <2.2e-16 | <2.2e-16 |
| R-HSA-597592 | Post-translational protein modification | 1425 | 594.55 | 1.2901 | <2.2e-16 | <2.2e-16 |
| R-HSA-8953854 | Metabolism of RNA | 673 | 280.79 | 1.6382 | <2.2e-16 | <2.2e-16 |
| R-HSA-8953897 | Cellular responses to external stimuli | 503 | 209.87 | 1.4819 | <2.2e-16 | <2.2e-16 |
| R-HSA-5663205 | Infectious disease | 382 | 159.38 | 1.5937 | <2.2e-16 | <2.2e-16 |
| R-HSA-72766 | Translation | 291 | 121.41 | 1.7955 | <2.2e-16 | <2.2e-16 |
| R-HSA-72203 | Processing of Capped Intron-Containing Pre-mRNA | 243 | 101.39 | 1.9135 | <2.2e-16 | <2.2e-16 |
| R-HSA-162906 | HIV Infection | 232 | 96.797 | 1.6839 | <2.2e-16 | <2.2e-16 |
| R-HSA-72172 | mRNA Splicing | 191 | 79.690 | 2.0329 | <2.2e-16 | <2.2e-16 |
| R-HSA-72163 | mRNA Splicing - Major Pathway | 183 | 76.353 | 2.0301 | <2.2e-16 | <2.2e-16 |
| R-HSA-6807070 | PTEN Regulation | 140 | 58.412 | 1.9174 | <2.2e-16 | <2.2e-16 |
| R-HSA-5368287 | Mitochondrial translation | 93 | 38.802 | 2.1391 | <2.2e-16 | <2.2e-16 |

Figure 11: Pathway enrichment analysis Tab under Results Panel
